# EEG microstate complexity for aiding early diagnosis of Alzheimer’s disease

**DOI:** 10.1101/833244

**Authors:** Luke Tait, Francesco Tamagnini, George Stothart, Edoardo Barvas, Chiara Monaldini, Roberto Frusciante, Mirco Volpini, Susanna Guttmann, Elizabeth Coulthard, Jon T Brown, Nina Kazanina, Marc Goodfellow

**Author notes:** Corresponding author Email address (Luke Tait).

## Abstract

**Introduction:** Electroencephalogram (EEG) is a potentially useful clinical tool for aiding diagnosis of Alzheimer’s disease (AD). We hypothesized we can increase the accuracy of EEG for aiding diagnosis of AD using microstates, which are epochs of quasi-stability at the millisecond scale.

**Methods:** EEG was collected from two independent cohorts of AD and control participants and a cohort of mild cognitive impairment (MCI) patients with four-year clinical follow-up. Microstates were analysed, including a novel measure of complexity.

**Results:** Microstate complexity significantly decreased in AD, and when combined with a spectral EEG measure, could classify AD with sensitivity and specificity >80%. These results were validated on an independent cohort and were also found to be generalizable to predict progression from MCI to AD. Additionally, microstates associated with the frontoparietal network were altered in AD.

**Discussion:** EEG has the potential to be a non-invasive functional biomarker that predicts progression from MCI to AD.

## 1. Background

Alzheimer’s disease (AD) is a neurological disorder in which progressive neurodegeneration and synaptic dysfunction result in impairments in a range of cognitive domains. Early diagnosis of AD has a wide range of clinical, social, and economic benefits (Leifer, 2003; Dubois et al., 2014; Alzheimer’s Association, 2018), but at present the definitive diagnosis of AD is made only post-mortem. Available tools for a diagnosis of probable AD in the clinic are mostly based on cognitive, biochemical, and neuroimaging markers (Agronin, 2014). While biochemical and neuroimaging techniques such as CSF or PET measurements of amyloid and tau have high sensitivity, large multi-centre studies have shown specificity as low as 72% for prodromal diagnosis (Mattsson et al., 2009). Since these molecular markers do not measure brain function, combining them with functional markers is likely to be useful for early diagnosis. Neuropsychological evaluation assesses brain function, is non-invasive and relatively inexpensive, but is time consuming, subject to cultural and personal bias, and is unsuitable for prodromal diagnosis of AD in cognitively homogeneous cohorts of mild cognitive impairment (MCI) patients. For an accurate and objective diagnosis, particularly at early stages, biological markers must be used. Biochemical and neuroimaging techniques such as CSF or PET measurements of amyloid and tau are invasive and/or not cost-effective, hence not easily viable for the screening of large populations at the prodromal or preclinical stages of the disease (Lee et al., 2017; Wittenberg et al., 2019).

Electroencephalography (EEG) is a non-invasive measure of neuronal electrical activity in the brain. EEG is a promising diagnostic tool for disorders of the central nervous system, since it is low cost, non-invasive and currently implemented in healthcare systems around the world for diagnosis of epilepsy (Smith, 2005). The first interest in using EEG as a diagnostic tool for AD was motivated by the associations between AD and epileptic activity (Friedman et al., 2012), but subsequent studies demonstrated limited success using epileptiform activity in the EEG to aid with the diagnosis of AD (Liedorp et al., 2010; Vossel et al., 2013; Cretin et al., 2016). For this reason, focus has since shifted to EEG signature activities not directly related to network hyperexcitability, such as resting state power spectral and functional connectivity analyses (Babiloni et al., 2016). At present, EEG testing is not routine in assessment of AD as the sensitivity and specificity is not sufficiently high, but commonly used EEG signatures of AD including spectral slowing and altered functional connectivity (Babiloni et al., 2016) are typically assessed on a time scale of the order seconds to minutes (Gudmundsson et al., 2007), assuming stationarity over these epochs. Using a combination of measures containing complementary information may achieve greater classification accuracy (Poil et al., 2013; Simpraga et al., 2017), therefore we hypothesise that combining these measures with analyses of non-stationarity within these epochs, novel information can be gained that can improve the ability of the EEG to classify AD (Dierks et al., 1997; Musaeus et al., 2019).

One such method is EEG microstate analysis, which involves studying the instantaneous topographic maps of the EEG (Michel and Koenig, 2018). Studies of EEG microstates have remarkably found the EEG to be comprised of only a small number of topographic classes (Michel and Koenig, 2018), such that the EEG remains stable in a given class for periods of the order tens or hundreds of milliseconds before rapidly transitioning to another class. These rapidly switching periods of quasi-stability are hypothesised to be the electrophysiological correlates of the rapid activation and inactivation of the brain’s resting state networks relating to different functions underpinning information processing (Khanna et al., 2015; Michel et al., 2001; Britz et al., 2010; Musso et al., 2010; Milz et al., 2016; Lehmann et al., 1998), earning microstates the nickname “atoms of thought” (Lehmann et al., 1998). Thus, EEG microstates are a viable platform for exploring the changes in brain dynamics on the millisecond scale that may underpin impaired information processing and cognitive dysfunction in AD; indeed, alterations to microstates have been observed in healthy development and aging (Koenig et al., 2002) and a range of neurological disorders including frontotemporal dementia, schizophrenia, and depression (Khanna et al., 2015). This makes EEG microstate analysis a suitable candidate for biomarkers at a faster temporal scale than power spectral or functional network analysis.

Much interest has been given to how properties such as duration, coverage, and topography of microstates are altered in neurological disorders (Khanna et al., 2015). Alterations to patterns of transitions between classes have also been reported in disease (Lehmann et al., 2005; Nishida et al., 2013), suggesting that studying transitioning behaviour of microstates may give further mechanistic insights into cognitive impairment in AD as well as increasing sensitivity of electrophysiological biomarkers. However, the ‘syntax analysis’ (Lehmann et al., 2005; Nishida et al., 2013) used to analyse transitioning behaviour in these studies assumes the next microstate depends only on the present microstate (Markovian) and probabilities of transitions do not change over time (stationary). Recent work has brought into question the validity of these assumptions (Van De Ville et al., 2010; von Wegner et al., 2017). Here, we present a novel analysis of the transitioning behaviour of EEG microstates, which does not rely on these assumptions, by applying the Lempel-Ziv complexity (LZC) algorithm (Lempel and Ziv, 1976) to microstate sequences. LZC counts the number of unique sub-sequences within a sequence. Microstate sequences with low LZC are repetitive, exhibiting a limited number of transitioning patterns. Conversely high LZC is suggestive of complex transitioning. Because this measure does not restrict the analysis to Markovian effects, we hypothesise microstate LZC could give new insights into alterations to information processing and transitioning between active networks in AD on the millisecond scale, and potentially act as a more sensitive EEG-based neurophysiological signature of AD.

Here we explore microstate statistics in AD based on 20 seconds of resting state EEG, and the potential clinical applications of microstate LZC, combined with a power spectral measure (Tait et al., 2019), for classification of early stage non-medicated AD patients. We then validate this classifier on an independent set of EEG recordings acquired with standard 19 electrode systems (used in clinical settings) in an independent sample of medicated AD patients from a distinct geographical location. Finally, since a key limitation of many past studies is the lack of longitudinal data, we explore the combination of microstates LZC and spectral power to predict the conversion of MCI to AD, based on an MCI cohort followed up longitudinally for up to 4 years.

## 2. Methods

### 2.1. Participants

Detailed descriptions of participant recruitment, inclusion criteria, diagnosis of probable AD, and analysis of demographics are given in Supplementary Material 1. AD patients (n=21, 8 male) and amnestic MCI patients (n=25, 16 male) were recruited from memory clinics in the South West of England (SWE) following clinical assessment on a consecutive incident patient basis. The diagnosis of AD was determined by clinical staff according to DSM-IV (American Psychiatric Association, 2000) and NINCDS-ADRDA guidelines (McKhann et al., 1984). Age matched healthy older adult (HOA) controls (n=26, 14 male) were recruited from the memory clinics’ volunteer panels; they had normal general health with no evidence of a dementing or other neuropsychological disorder, according to NINCDS-ADRDA guidelines (McKhann et al., 1984). MCI participants who did not receive a dementia diagnosis within the four years following data acquisition were classified as stable (MCIs, n=7, 5 male), those who had received an AD diagnosis were classified as converters (MCIc, n=4, 4 male), and the remaining patients were excluded from the analysis (see Supplementary Material 1). All participants were free from medication known to affect cognition. Cognitive status at the time of data acquisition was quantified using the mini-mental state examination (MMSE) (Folstein et al., 1975).

A test cohort of AD patients (n=9, 3 male) and HOA controls (n=7, 4 male) was used to test generalizability of EEG biomarkers. Patients were recruited on a consecutive incident basis from the San Marino data-base of dementia. The diagnosis of AD was determined according to the IWG-2 criteria (Dubois et al., 2014) by neurologists at the Republic of San Marino State Hospital. HOA controls were recruited from caregivers of patients. In addition, all the AD patients from the San Marino cohort were being administered anticholinesterase drugs. Episodic memory impairment was quantified using the Rey Auditory Verbal Learning Test (RAVLT) (Carlesimo et al., 1996).

All participants provided written informed consent before participating and were free to withdraw at any time. All procedures were approved by relevant ethics committees (see Supplementary Material 1).

### 2.2. EEG Collection and Pre-processing

20 seconds of resting-state, eyes-open EEG was used for all subjects. Data was sampled at 1000/512 Hz and recorded with 64/19 channels (SWE/RSM cohorts respectively). Data from both cohorts was preprocessed identically following previously described steps (Tait et al., 2019). Details of the 20 second epoch selection and pre-processing can be found in Supplementary Material 2.

### 2.3. Microstate Analysis

#### 2.3.1. Microstate Extraction

Microstates were extracted using a k-means clustering method based on that of Koenig et al. (1999), using a k-means++ algorithm used to select the initial k maps (Arthur and Vassilvitskii, 2007), and 20 repetitions Koenig et al. (1999). The Krzanowski-Lai criterion was used to assess the optimum number of microstates (Hatz et al., 2015). The median optimum over all subjects in the training set was *k* = 4, with no differences between groups. To ensure maps were comparable within cohorts (HOA or AD), a global clustering algorithm was performed (Khanna et al., 2014) for each cohort, and global maps between cohorts were aligned by calculating correlation coefficients of the maps and visual inspection. Additional details can be found in Supplementary Material 2.

### 2.4. Analysis of microstates in Alzheimer’s disease

#### 2.4.1. Basic Microstate Statistics

Mean microstate duration for each class and percent time spent within a class (coverage) were calculated (Khanna et al., 2015). Analysis of microstate syntax, i.e. Markovian transitioning, was performed by extracting the transition matrix from the data by counting the number of transitions from class *i* to class *j* and normalizing by the total number of transitions (Lehmann et al., 2005; Nishida et al., 2013). For all measures, a two-way ANOVA was performed to assess statistical differences. Since ANOVA tests are parametric, significant results were post-hoc validated using non-parametric Wilcoxon rank sum tests. Topographic differences between groups were assessed with class-wise topographic analyses of variance (TANOVA) (Milz et al., 2016) (Supplementary Material 2). The minimum possible P-value was *P* = .001, deemed sufficient to identify significant differences in topographies.

#### 2.4.2. Microstate Complexity

Here we present a novel measure of EEG microstate transitioning that, unlike the Markovian syntax analysis, does not rely on assumptions of stationarity or Markovianity (von Wegner et al., 2017). The measure, referred to as *C*, involves calculating the Lempel-Ziv complexity (LZC) (Lempel and Ziv, 1976) of the microstate transitioning sequence. The LZC of a string is defined as the number of different substrings within the string when read from left to right. A string is said to have low complexity if there are a small number of frequently repeating sequences. The algorithm for computing *C* is outlined in Supplementary Fig. S1 and Supplementary Material 3. LZC has in the past been used to calculate the complexity of a univariate EEG signal (Supplementary Material 3), but we believe use of this measure to explore the complexity of the microstate sequences is novel. All statistical comparisons of microstate LZC between groups was performed using non-parametric Wilcoxon rank sum tests.

## 3. Results

### 3.1. Participant Demographics

Participant demographics, including age and neuropsychological scores, are outlined and analysed in Table 1, Supplementary Material 1 and Supplementary Table S1-S2. In both cohorts, AD patients demonstrated reduced neuropsychological test scores compared to healthy controls. There were no differences in neuropsychological test score between MCI stable (MCIs) and MCI-to-AD converter (MCIc) at baseline.

**Table 1:**
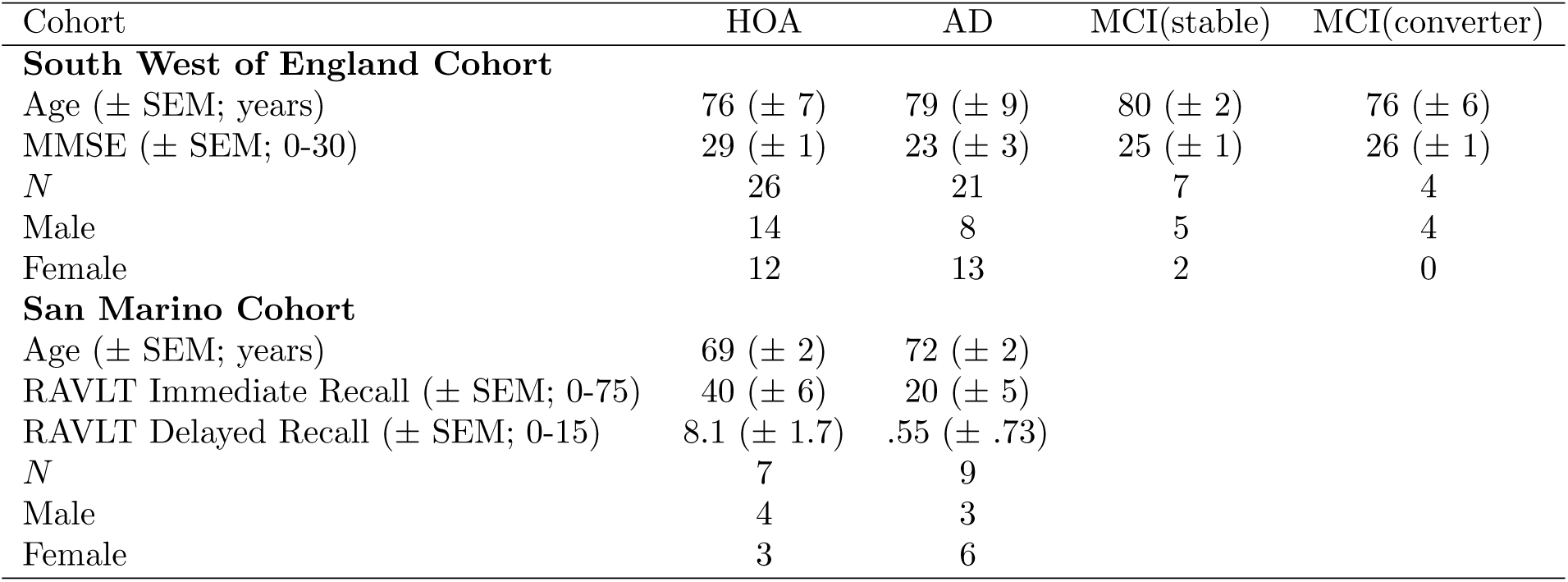
Data for healthy older adults and Alzheimer’s disease subjects for the training and test cohorts.

| Cohort | HOA | AD | MCI(stable) | MCI(converter) |
| --- | --- | --- | --- | --- |
| **South West of England Cohort** |  |  |  |  |
| Age (± SEM; years) | 76 (± 7) | 79 (± 9) | 80 (± 2) | 76 (± 6) |
| MMSE (± SEM; 0-30) | 29 (± 1) | 23 (± 3) | 25 (± 1) | 26 (± 1) |
| *N* | 26 | 21 | 7 | 4 |
| Male | 14 | 8 | 5 | 4 |
| Female | 12 | 13 | 2 | 0 |
| **San Marino Cohort** |  |  |  |  |
| Age (± SEM; years) | 69 (± 2) | 72 (± 2) | | |
| RAVLT Immediate Recall (± SEM; 0-75) | 40 (± 6) | 20 (± 5) | | |
| RAVLT Delayed Recall (± SEM; 0-15) | 8.1 (± 1.7) | .55 (± .73) | | |
| *N* | 7 | 9 |  |  |
| Male | 4 | 3 |  |  |
| Female | 3 | 6 |  |  |

### 3.2. Analysis of microstates in Alzheimer’s disease

Topographies for each of the four microstate classes in the SWE HOA and AD data are shown in Figure 1, and in the HOA subjects largely align with the classical microstate classes A-D associated with wakeful rest (Khanna et al., 2015; Michel and Koenig, 2018), which are electrophysiological correlates of the auditory (A), visual (B), saliency (C), and frontopariental working memory/attention (D) resting state networks (Michel and Koenig, 2018; Britz et al., 2010). Class D was significantly altered in AD (*P* = .001; Figure 1D). eLORETA was used to explore cortical generators underpinning this alteration (Supplementary Material 2). Figure 2A shows the eLORETA solution of the instantaneous map given by the difference of the global class D maps for HOA and AD. We subsequently calculated the eLORETA solution on a subject wise basis and calculated voxel-wise *t*-statistics to quantify spatially distributed differences in AD. Parietal sources were less activated in the AD subjects, particularly weighted more towards the left hemisphere (Figure 2B-C).

**Figure 1:**
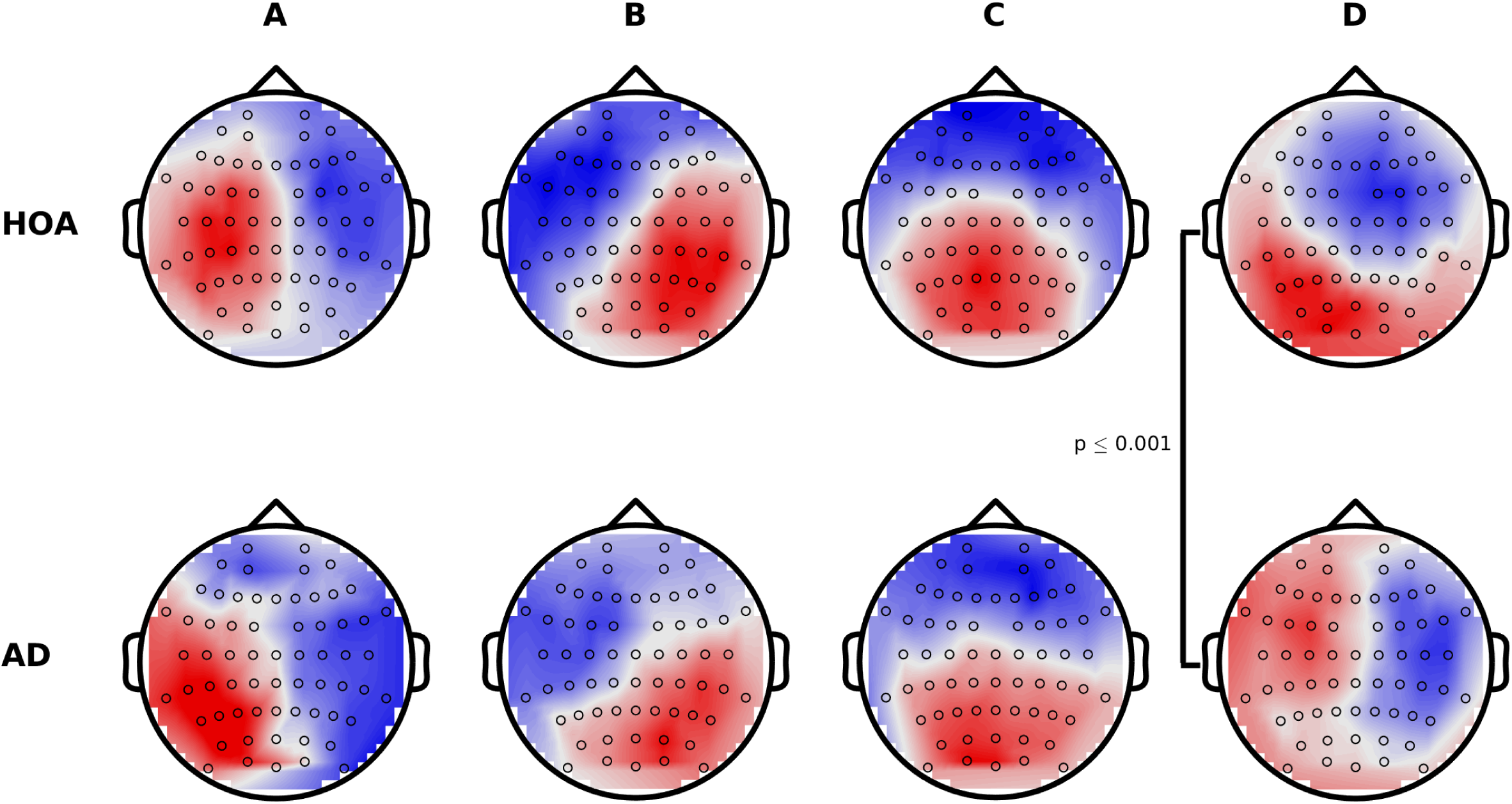
Microstate topographies for the four classes. (Top) Globally clustered maps for HOA cohort, for classes A-D from left to right. (Bottom) As above, but for the AD cohort. Black circles mark the electrode locations.

**Figure 2:**
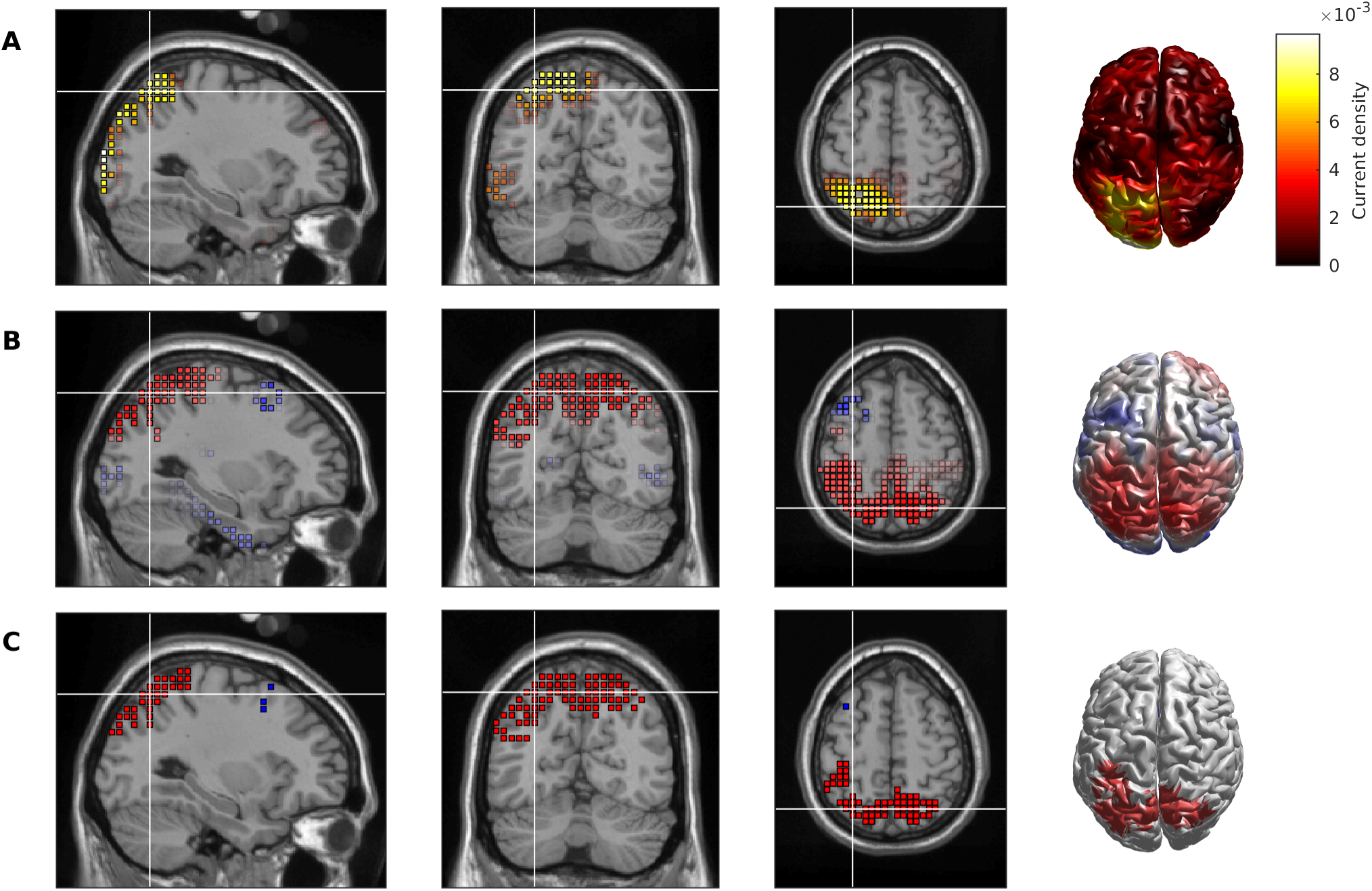
Cortical source generators underpinning alterations to microstate class D in AD. (A) Absolute value of the eLORETA solution to the instantaneous topography given by taking the difference between the global class D maps for HOA and AD. (B) *t*-statistic for voxel-wise comparisons of the subject-wise class D maps for HOA vs AD. Red indicates absolute value of current density is larger for HOA than AD, whilst blue is AD > HOA. (C) Voxels with *t*-values such that *P* < .05. Red voxels indicate HOA > AD, and blue voxels are AD > HOA.

Having found changes to the topography and underlying cortical generators to class D in AD, we next explored whether the transitioning behaviour of the microstate sequences were altered. For each subject and microstate class, mean duration, coverage, and the Markovian syntax transition matrix were extracted. Supplementary Tables S3-S5 show the ANOVA tables for these tests. For mean duration, there was a significant disease group term (*P* = .0019), suggesting a significant increase in mean microstate duration in AD, averaged over all classes. We verified this significant increase in mean microstate duration with a non-parametric Wilcoxon rank sum (*P* = .0214, Figure 3A). No significant results were found for coverage of microstates or Markovian transitioning. All analyses were additionally conducted on a pairwise basis for class D to further verify that changes to the topography of this class in AD did not alter transitioning statistics. No significant differences were found.

**Figure 3:**
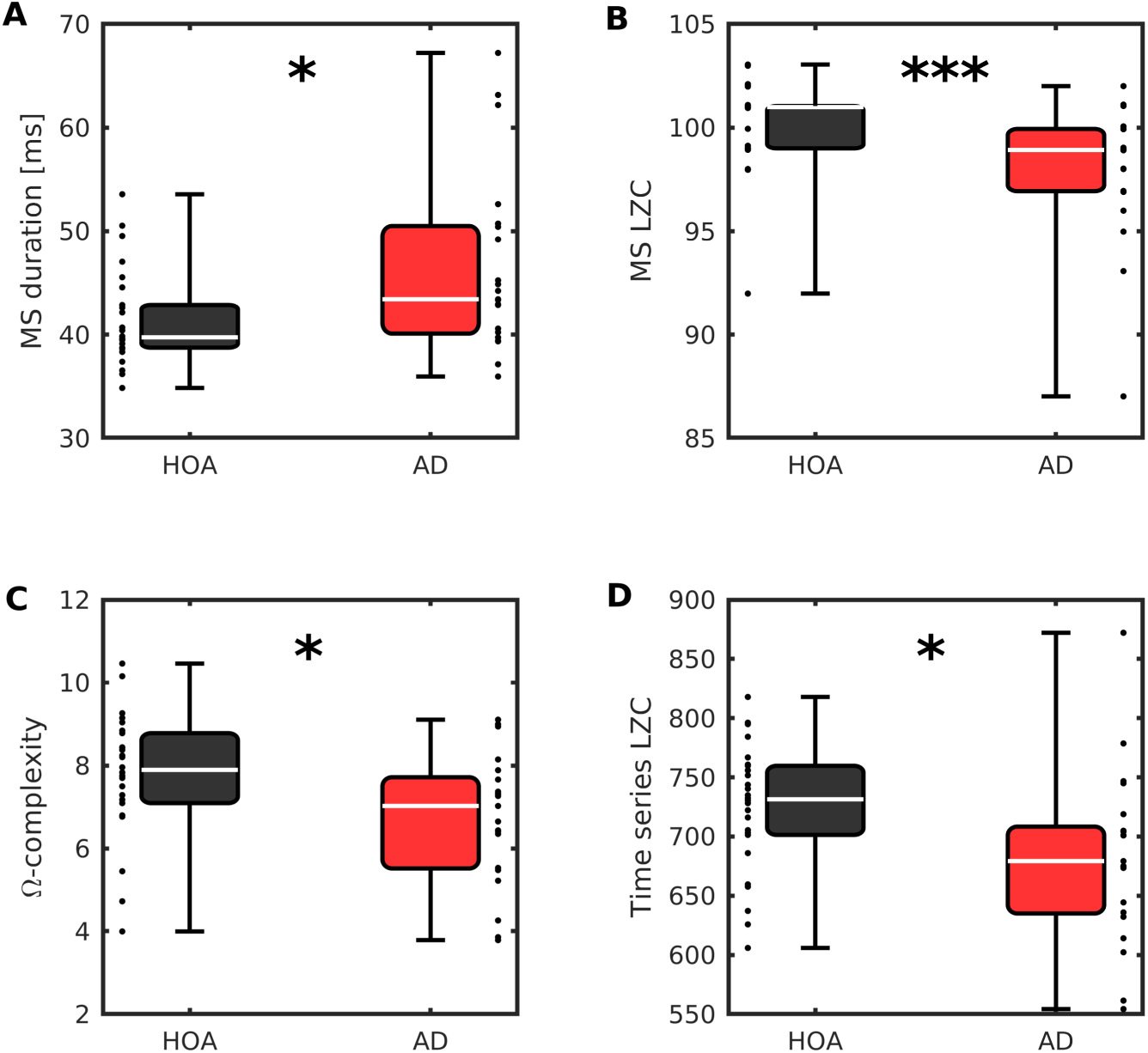
Microstate and complexity statistics are significantly altered in AD. (A) Mean duration of microstates. (B) Microstate LZC. (C) Ω-complexity. (D) Time series LZC. Descriptions of classical complexity measures C-D are given in Supplementary Material 3. Stars denote effect size of Mann-Whitney U test: * *P* < .05, ** *P* < .01, *** *P* < .001. Points next to boxplots show values for each participant.

Studies have suggested that microstate transitioning is non-Markovian (Van De Ville et al., 2010; von Wegner et al., 2017). We therefore additionally calculated the Lempel-Ziv complexity (LZC) of the microstate transitioning sequence (*C*). *C* was significantly reduced in AD compared to HOA (*P* = .0023; Figure 3B), suggesting that higher order or non-Markovian alterations to microstate transitioning are present in the EEG of AD patients, to which a Markovian syntax analysis is not sensitive. Additionally, this effect size was notably larger than other classical measures of EEG complexity used in past AD literature (Figure 3C-D, Supplementary Material 3).

### 3.3. Microstate complexity, combined with a single spectral feature, is useful as a robust and generalizable biomarker of AD

Spectral slowing of the EEG in AD has been widely reported (Babiloni et al., 2016), and has been identified as a powerful tool for classification of AD from the EEG (Poil et al., 2013; Simpraga et al., 2017). A key aim of this study was to develop EEG microstate biomarkers for aiding diagnosis of AD, since we hypothesised that the difference in timescale between microstate measures and spectral measures would result in orthogonal information to increase classifier accuracy. Microstate LZC has a large effect size separating people with AD from controls and is uncorrelated with spectral slowing (Supplementary Material 3), so we next combined microstate LZC (*C*) with theta relative power (*θ*RP), a proxy for slowing previously identified in this dataset (Tait et al., 2019) in a support vector machine (SVM) classifier (Supplementary Material 2).

Table 2 shows the results of 10-fold cross validation of the classifier, trained on the 64 channel SWE EEG analysed above. When features were combined, classification rate (CR), sensitivity, and specificity were all greater than 80%, with a CR of 85.1%, demonstrating a notable improvement over the use of these features independently. Figure 4A shows this classifier in 2D (*θ*RP, *C*) space.

**Table 2:**
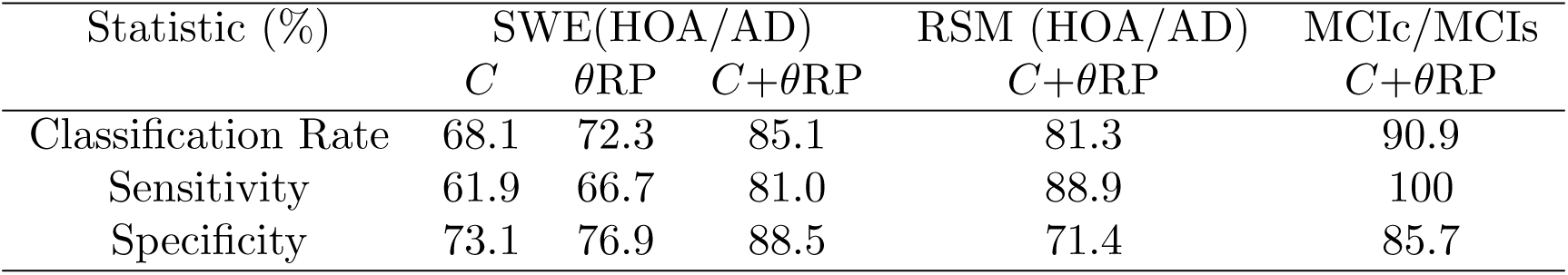
Classification statistics from EEG measures in the training and test set.

**Figure 4:**
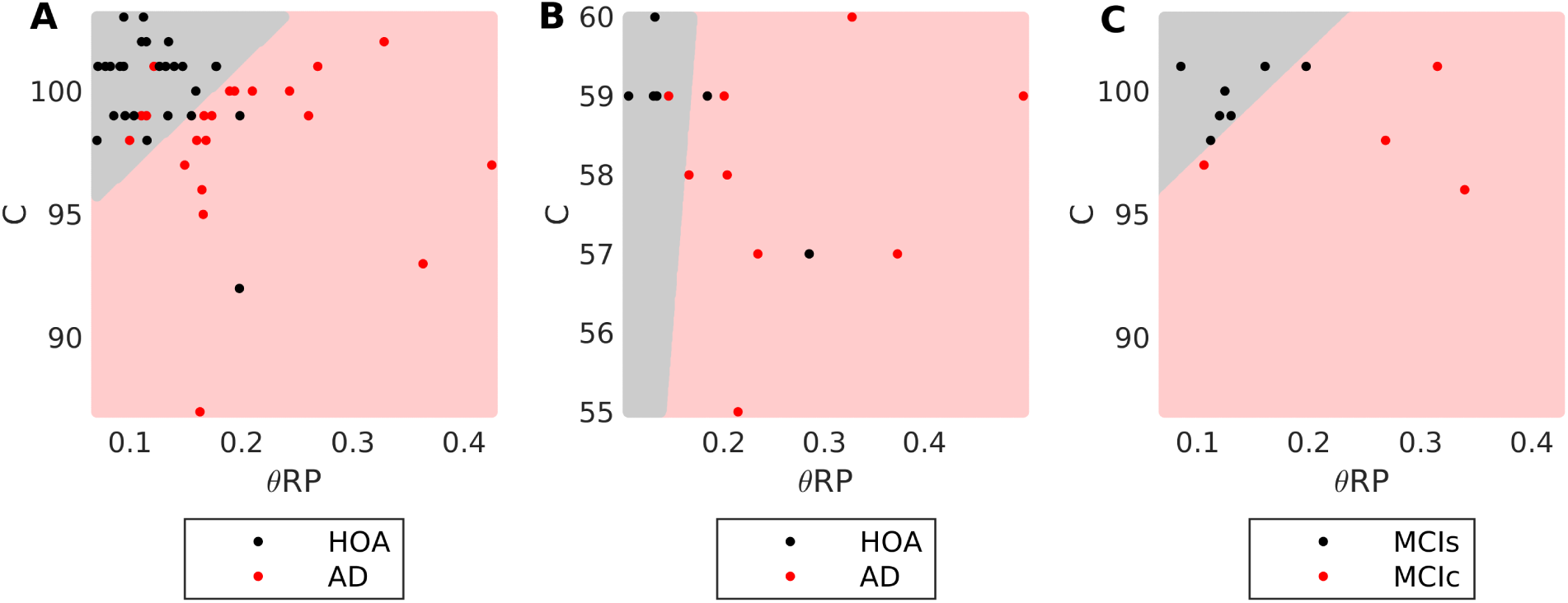
Separation of AD, HOA, and MCI using a SVM *θ*RP+*C* classifier. The SVM predictor classifies points within the pale red region as AD and points within the gray region as HOA. (A) Training data set overlaid on 64 channel model. (B) RSM data set overlaid on 19 channel classifier model. (C) MCI data set overlaid on 64 channel model. All models were trained on the data set shown in A. Blue circles in A show the support vectors for training in the 64 channel model.

To validate the classifier, the RSM cohort of clinical EEG was used as an independent test set. The SWE training data was spatially and temporally down-sampled to the same format as the RSM EEG, and the *θ*RP+*C* classifier re-trained on this down-sampled data set. Table 2 and Figure 4B show the results of the classifier on the independent RSM test data. The CR of the test data was 81.3% (13/16 subjects), with sensitivity 88.9% (8/9 AD patients classified as AD by the model) and specificity 71.4% (5/7 HOA subjects classified as HOA by the model). These results suggest the SVM classifier is generalizable to new clinical data sets and is not due to overfitting to our data.

### 3.4. Microstate complexity and slowing have predictive power for diagonsing prodromal AD

Finally, to test whether the classifier presented here is useful as a tool for prodromal diagnosis of AD (Dubois et al., 2014), i.e. predicting whether mild cognitive impairment (MCI) is due to prodromal AD, EEG recorded from a test set of MCI patients was classified by the model. Subjects were classified as MCIs or MCIc based on their clinical diagnosis four years following data acquisition. At the time of data acquisition, there was no difference in clinical diagnosis or MMSE scores for the subjects (Table 1, subsection 3.1). Results of the classification are shown in Figure 4C and Table 2. The model had a classification rate of 90.9%, correctly predicting the four year diagnosis of 10/11 MCI patients. One MCIs subject was incorrectly classified as a converter.

## 4. Discussion

The aim of this study was to validate microstate analysis as a predictive tool for the diagnosis of probable AD. A classifier was trained on a cohort of people with AD and HOA recruited from the South West of England (SWE), using 64-channel EEG. Subsequently, the classifier was validated on an independent cohort recruited from the Republic of San Marino (RSM), using a standard 19-channel EEG machine for clinical use. Whilst the RSM AD population was treated with acetylcholinesterase (AChE) inhibitors, the SWE cohort was unmedicated. This is an important fact in support of this method for early diagnosis, as the use of AChE inhibitors to treat people with AD is very common and a possible confounding factor for functional measures. The accuracy of our method was measured against a clinical diagnosis of probable AD based on IWG-2 (Dubois et al., 2014) or DSM-IV (American Psychiatric Association, 2000) and NINCDS-ADRDA guidelines (McKhann et al., 1984).

### 4.1. Microstate complexity measure

In this work we present a novel application of Lempel-Ziv complexity (LZC) (Lempel and Ziv, 1976) to microstate EEG sequences for the diagnosis of AD. LZC was chosen to study microstate transitions over classical Markovian syntax analysis since microstate transitioning is neither Markovian nor stationary (Van De Ville et al., 2010; von Wegner et al., 2017). We found that the LZC of microstate transition sequences had a notably larger effect size in separating AD patients from controls than Markovian syntax analysis, suggesting higher order alterations to transitioning between microstates in AD.

As a multivariate measure, microstate LZC additionally captures both spatial and temporal complexity, resulting in a larger effect size than previously reported measures of EEG complexity (Supplementary Material 3). A closely related univariate measure of EEG complexity is time series LZC, which has in the past been used to identify reduced complexity of the EEG in patients with AD by binarizing a univariate EEG time series based on a threshold and then calculating the LZC of this binary sequence (Supplementary Material 3). Application of LZC to the microstate sequences is an improvement over this method for several reasons. By using microstates, binning of the EEG is chosen based on repeating spatiotemporal patterns which have some neurophysiological basis related to active networks (Michel and Koenig, 2018; Milz et al., 2016), as opposed to an arbitrary threshold. Furthermore, microstate LZC accounts for the multivariate nature of the EEG, not accounted for in classical time series LZC. Microstate LZC is a spatially extended version of the LZC method classically used in EEG literature. Finally, unlike time series LZC (Dauwels et al., 2011), microstate LZC is uncorrelated with EEG slowing (Supplementary Material 3), meaning it is a useful EEG biomarker of AD when combined with slowing.

### 4.2. Alterations to class D and the frontoparietal network

The topography of microstate class D was altered in AD. Nishida et al. (2013) studied microstate topographies in AD and found no alterations to any of the four microstate classes. Differences in the number of channels is unlikely to explain these inconsistencies in results, as microstates are reliable with 8 or more channels (Khanna et al., 2014). The use of eyes-open data is also an unlikely to give inconsistent results (Seitzman et al., 2017) (Supplementary Material 4). However, methodological differences may potentially explain opposing findings; in Nishida et al. (2013) all participants were used for defining the topographies of the four classes, whilst here classes were defined independently for each cohort.

It has been suggested that class D is related to the frontoparietal network and the attention and working memory cognitive domains (Khanna et al., 2015; Britz et al., 2010; Nishida et al., 2013; Seitzman et al., 2017). Reduced parietal activation underpinned the change in topography of class D in AD, supporting the hypothesis of dysfunction of the frontoparietal network. A possible mechanism for this is disrupted frontoparietal white matter integrity, since altered frontoparietal functional and effective connectivity have been reported in recent fMRI studies of AD (Neufang et al., 2011; Zhao et al., 2018).

### 4.3. Alterations to microstate duration and transitioning statistics

In this study, mean microstate duration was found to increase in AD. Reports of altered microstate duration are inconsistent in the past literature, which has found increased (Ihl et al., 1993), decreased (Dierks et al., 1997; Strik et al., 1997; Stevens and Kircher, 1998), or unchanged (Musaeus et al., 2019; Nishida et al., 2013) durations in AD. Of these studies, the two which have used modern clustering methods comparable to this study were those that identified no differences in duration. Eyes-open vs eyes-closed EEG affects microstate duration (Supplementary Material 4) (Seitzman et al., 2017), potentially explaining our conflicting results. Interestingly, we might expect microstate duration to increase in AD due to slowing of the EEG (von Wegner et al., 2017) which is well established in AD (Babiloni et al., 2016) - this was the original hypothesis made by Dierks et al. (1997), who expressed their surprise that a decreased microstate duration was found instead. Increased duration and reduced microstate LZC are suggestive of slower and more repetitive transitioning between active networks in AD, possibly indicating less complex information processing and giving insight into the mechanisms underpinning cognitive deficits in AD.

### 4.4. Conclusions

In this work we validated microstate analysis, in combination with power spectral analysis, as a low-cost, non-invasive tool for aiding the diagnosis of AD. Additionally, our observations can provide crucial insight into the mechanisms underpinning cognitive impairment in AD. Alterations to the frontoparietal network (namely parietal inactivation) was shown to relate to a changing topography in microstate class D. This network and microstate class are related to attention and working memory (Britz et al., 2010; Nishida et al., 2013; Seitzman et al., 2017), which are impaired early in the AD staging (Perry and Hodges, 1999). Microstate duration was found to increase in AD, whilst a novel application of Lempel-Ziv complexity to the microstate transitioning found decreased complexity in AD. These results are suggestive of slower and more repetitive microstate transitions, which potentially reflects similar attributes to the transitioning between active brain networks associated with a range of cognitive domains (Khanna et al., 2015; Michel and Koenig, 2018; Britz et al., 2010; Musso et al., 2010; Milz et al., 2016; Lehmann et al., 1998). Preliminary data suggest use of microstate complexity as a biomarker can aid with early diagnosis and prediction of future conversion to AD. Whilst medication is known to alter microstate statistics (Lehmann et al., 1993; Yoshimura et al., 2007), our classifier accurately provided an AD diagnosis in independent, geographically distant cohorts, one of which was unmedicated whilst the other was treated with AChE inhibitors. The mechanistic insights presented in this study, paired to future, preclinical characterization of cellular correlates of microstate alterations in transgenic rodent models of AD pathologies may aid future drug development and more accurate diagnostic and prognostic tools for AD.

## Acknowledgements

This work was supported by the EPSRC [grant numbers EP/P021417/1 and EP/N014391/1] (MG); a Wellcome Trust Institutional Strategic Support Award (https://wellcome.ac.uk/) [grant number WT105618MA] (MG); the Alzheimer’s Society in partnership with the Garfield Weston Foundation [grant reference 231] (LT); University Research Fellowship from the University of Bristol (NK). Work by EB was jointly supported by the University of San Marino and ISS. The funders had no role in study design, data collection and analysis, decision to publish, or preparation of the manuscript.

