## Supplementary Material for "EEG microstate complexity for aiding early diagnosis of Alzheimer’s disease"

#### 1. Participant details and demographics

##### *1.1. Training data (SWE HOA/AD)*

AD patients (*n* = 21, 13 female, 8 male) were recruited from memory clinics in the South West of England (SWE) following clinical assessment on a consecutive incident patient basis. The diagnosis of AD was determined by clinical staff using neurological, neuroimaging, physical and biochemical examination together with the results of family interview, neuropsychological and daily living skills assessment according to DSM-IV [1] and NINCDS-ADRDA guidelines [2]. Age matched healthy older adult (HOA) controls (*n* = 26, 12 female, 14 male) were recruited from the memory clinics' volunteer panels; they had normal general health with no evidence of a dementing or other neuropsychological disorder, according to NINCDS-ADRDA guidelines [2]. All participants were free from medication known to affect cognition (e.g. cholinergic medications prescribed to treat dementia symptoms, anti-psychotics, anti-depressants, benzodiazepines, Warfarine, etc) and had no history of transient ischemic attack, stroke, significant head injury, psychiatric order, or neurological disease with non-AD aetiology. Cognitive status was quantified using the mini-mental state examination (MMSE) [3]. Participants provided written informed consent before participating and were free to withdraw at any time. All procedures for this cohort were approved by the National Research Ethics Service Committee South West Bristol (Ref. 09/H0106/90).

##### *1.2. Test data (RSM HOA/AD)*

A test cohort of patients was used to test generalizability of EEG biomarkers. All participants were 45-85 years old. AD patients in the test cohort (*n* = 9, 6 female, 3 male) were recruited on a consecutive incident basis from the San Marino data-base of dementia. A diagnosis of probable AD was defined according to the IWG-2 criteria [4] by neurologists at the Republic of San Marino State Hospital: briefly, volunteers were included in the study if they showed an objective evidence of an amnestic syndrome of the hippocampal type (based on significantly impaired performance on an episodic memory test with established specificity for AD) and if they showed *in vivo* evidence of Alzheimer's pathology (decreased $A\beta_{1-42}$ together with increased T-tau and P-tau in CSF). In addition, all the AD patients from the San Marino cohort were being administered anticholinesterase drugs, in accordance with the Good Clinical Practice regulations of the Republic of San Marino. Age matched healthy controls (*n* = 7, 4 male, 3 female) were recruited from caregivers of patients. All subjects were free from cognitive disorders with a non-AD aetiology, sensory disorders, and psychiatric disorders. Participants provided written informed consent before participating and were free to withdraw at any time. All procedures for this cohort were approved by the Republic of San Marino Ethical Committee for Research and Experimentation (Ref. 0015 SM) and the University of Exeter Medical School Research Ethics Committee.

##### *1.3. MCI data (SWE MCIs/MCIc)*

In addition to the two cohorts of AD patients and healthy controls, a cohort of amnestic mild cognitive impairment (MCI) patients (*n* = 25) were also recruited from memory clinics in the South West of England under the guidelines for inclusion and procedure approvals of the SWE HOA/AD cohort. The aim of this cohort was to test the ability of EEG biomarkers to aid in diagnosis of prodromal AD, i.e. MCI with an AD aetiology [4], therefore dementia status was reassessed four years following data acquisition and patients who did not receive an AD diagnosis at follow-up (*n* = 7, 5 male, 2 female) were classified as MCI-stable (MCIs), whilst patients who had a diagnosis of AD at the four year follow-up (*n* = 4, 4 male) were classified as prodromal AD or MCI-converters (MCIc). The remaining 14 patients were either unavailable, deceased, or had a diagnosis of non-AD or mixed dementia at the four year follow-up and were excluded from the analysis.

#### 1.4. *Demographics*

Participant demographics are outlined in Table 1 of the main text. Across the six cohorts (SWE/RSM HOA/AD, MCIs, MCIc), there was a significant effect of age ($\chi^2 = 12.63$, $P = .0271$, Kruskal-Wallis test). Pairwise Mann-Whitney U tests demonstrated that the ages of the RSM HOA cohort were different to the SWE HOA, SWE AD, and MCIs cohorts at a level of $P < .05$. Additionally, RSM AD cohort was different in age to SWE AD and MCIs at a level of $P < .05$. However, following correction for multiple hypotheses, no differences were significant. The *P*-values for all pairwise comparisons of age are given in Supplementary Table S1.

Mini-mental state examination scores were available for the SWE cohorts. Across these four cohorts, there was a significant effect of MMSE score ($\chi^2 = 37.35$, $P = 3.89 \times 10^{-8}$). Pairwise testing found that, with the exception of MCI stable vs converters ($p = 0.7939$, Mann Whitney U test), there were differences in MMSE between all pairs of cohorts to $p < 0.05$. These results remained significant following false discovery rate correction for multiple hypotheses. All *P*-values for pairwise comparisons of MMSE are given in Supplementary Table 2.

The assessment of the cognitive state of the RSM cohort was based on a different battery of tests. The results are summarised in Supplementary Table S3.

### 2. Methods for EEG acquisition, pre-processing, and analysis

#### 2.1. *Acquisition and pre-processing*

Resting-state, eyes open EEG was recorded from all subjects. For the participants recruited from the South West of England (SWE), data was sampled at 1 kHz and recorded with 64 channels. A single 20s epoch of EEG was chosen from a period prior to the beginning of a battery of cognitive tasks for each subject. This period, and the subsequent battery of tasks, were consistent across the two groups. For participants recruited from the Republic of San Marino (RSM), data was sampled at 512 Hz and recorded with 19 channels in the 10-20 format. For two subjects, data was sampled at 128 Hz and linearly interpolated to 512 Hz. TAPEEG was used to automatically detect optimal 20s epochs of EEG [5]. Data from both cohorts was pre-processed identically. Visual and cardiac artefacts were removed using independent component analysis, whilst data was bandpass filtered (1-200 Hz), demeaned, detrended, and re-referenced to average. These pre-processing steps were performed using the Fieldtrip toolbox for EEG/MEG-analysis, developed at the Donders Institute for Brain, Cognition and Behaviour [6] (http://www.ru.nl/neuroimaging/fieldtrip).

#### 2.2. *Cortical Source Localization using eLORETA*

Cortical source localization was performed using the eLORETA algorithm [7,8], which has been used in recent literature to source localize EEG microstates [9–11], implemented in the sLORETA/eLORETA software package (http://www.uzh.ch/keyinst/loreta.htm). This package uses a 3 layer BEM head model [12] based on the Montreal Neurological Institute average MRI brain map (MNI152) [13] and a 6239 voxel source space limited to a cortical grey matter surface. This model has been validated and demonstrated to perform to a similar standard as individual MRI derived head models [12]. Visualization of results was performed using custom written Matlab routines and used the 'Colin27 2016' template MRI and cortical surface implemented in Brainstorm [14], which is documented and freely available for download online under the GNU general public license (http://neuroimage.usc.edu/brainstorm). To compare two topographic maps, for example the cluster average for a class between groups, the difference between maps was computed and then source localized using eLORETA. The source space representation of the topographic difference map shows the difference between cortical generators underpinning the two maps, since the forward model assumes linear mixing, i.e. $Y = \Phi X$ therefore $Y_2 - Y_1 = \Phi(X_2 - X_1)$, where $Y$ is the topographic map in sensor space, Φ is the leadfield, and $X$ is the set of cortical sources generating the map. To gain a measure of statistical effect size between groups, the subject-wise maps are source localized and then a $t$-statistic computed on a voxel-wise basis [10].

#### 2.3. *Microstate extraction*

Microstates were extracted using a $k$-means clustering method based on that of Koenig et al [15]. The global field potential (GFP) of the EEG at each time point is given by the standard deviation over all electrodes at that time point [15]. Topographic maps at the peaks of the GFP were extracted as these have the highest signal-to-noise ratio [16]. GFP peak maps were then clustered using the $k$-means clustering algorithm described by Koenig et al. [15], with a $k$-means++ algorithm used to select the initial $k$ maps [17]. For a given value of $k$, the clustering algorithm was repeated 20 times and the resulting cluster maps that explained most variance were chosen to be the optimum [15]. Microstates were defined to switch at the midpoints between GFP peaks, and all time points within a microstate were assigned to the class of the GFP peak.

The choice of $k$ is non-trivial. For each subject, we therefore used the Krzanowski-Lai criterion [18] to assess the optimum number of microstates [19]. In this data, there were no significant differences between the optimum number of microstates in AD and HOA as identified by a non-parametric Wilcoxon rank sum test. The median optimum over all subjects was four, so $k = 4$ maps were used for all subjects.

Since $k$-means clustering is performed on a subject-wise basis, it is important for statistical purposes that the four classes are comparable both within and between cohorts. Firstly, for each cohort (HOA or AD), a global clustering algorithm [20] was performed to relabel classes such that classes were comparable between subjects within a cohort. Secondly, visual inspection and calculation of correlation coefficients of the four globally clustered centroid maps per cohort were used to align classes such that class labels were comparable between cohorts.

#### 2.4. *Topographic Analysis of Variance (TANOVA)*

Topographic differences between groups were assessed with class-wise topographic analyses of variance (TANOVA) [10]. The topographic dissimilarity between the HOA and AD group average maps for a given class was calculated as the GFP of the difference between the maps. Group labels were subsequently permuted 999 times and the topographic dissimilarity recalculated to generate a non-parametric surrogate null distribution of dissimilarities. TANOVA $P$-values were computed vs this null distribution by rank ordering the dissimilarities of the surrogates and original data in descending order, then computing the position of the original data set in this distribution. Finally, this was divided by 1000 (999 surrogates + 1 original data set). Therefore a minimum possible $P$-value of .001 was available, deemed sufficient to identify significant differences in topographies.

#### 2.5. *Classification of AD*

Classification was performed through use of a binary support vector machine [21] implemented in Matlab's function 'fitcsvm' (https://uk.mathworks.com/help/stats/fitcsvm. html). The box constraint and kernel scale were optimized, whilst all other parameters were set to default. The training set (SWE HOA/AD) was used to build an SVM model to classify AD status of participants. All classification rates, sensitivities, and specificities reported for the training set were 10-fold cross validated to avoid overfitting. The model, trained to the SWE HOA/AD training set, was tested on the RSM HOA/AD and SWE MCIs/MCIc test data sets. When testing the model on the former test set, the training data was first spatially and temporally downsampled to the same 19 channels and 512 Hz as the test data and the classifier was re-trained.

### 3. Microstate Complexity

#### *3.1. Calculation of microstate LZC*

In this manuscript, a novel measure of microstate transitioning complexity was presented. The measure involves applying the Lempel-Ziv complexity (LZC) algorithm [22] to the EEG microstate sequences. LZC is defined as the number distinct sub-sequences within a sequence when read left to right, and was originally developed as the basis of a number of lossless compression algorithms that work by replacing repeated sub-sequences within markers [23,24]. The algorithm for computing LZC is described in Supplementary Figure S1. A string is said to have low LZC if there are a small number of frequently repeating sequences, i.e. if the string can be compressed to a very small amount of data. Microstate sequences with low LZC are therefore repetitive, with a limited number of transitioning patterns within the sequence, whilst high LZC is suggestive of complex and varied transitioning.

To calculate microstate LZC, the continuous multivariate EEG was first converted to a discrete sequence by computing the microstate sequence for the data; each time point was mapped from $\mathbb{R}^{64}$ to a single, discrete value in the set of microstate classes $\{A,B,C,D\}$, resulting in a single string of length equal to that of the number of sampling points in the original EEG time series. The sequence was then reduced to its transitioning sequence (e.g. a sequence AAABBCCCDAADD would be reduced to ABCDAD). Finally, the LZC of this sequence was then computed. The choice of computing LZC on the transitioning sequence instead of the raw microstate sequence was motivated by the fact the latter is strongly correlated with slowing of neuronal oscillations while the former is not (LZC of transitioning sequence vs $\vartheta$RP [25], $r$=-0.2734, $P$=.1260; LZC of raw microstate sequence vs $\vartheta$RP, $r$=-0.6198, $P=6.8\times10^{-6}$). Since LZC tends to increase with sequence length, only the only first $N$ entries of the switching-sequence were used to calculate microstate LZC. Here, we chose $N$ = 250 as this number is small enough that all subjects had a transitioning sequence of length greater than or equal to this number.

#### *3.2. Comparison with classical EEG complexity measures*

To compare how our novel measure of EEG complexity based on the LZC of microstate sequences performs compared to classical measures of EEG complexity, we additionally computed two such measures often used in the literature. Firstly, past literature has used an alternate method for calculation of the LZC [26–31], which we label time-series LZC ($C^{TS}$). For a given EEG time series (i.e. a single electrode), each time point is mapped from $\mathbb{R}$ to a value from a discrete set, by binarizing the data based on an arbitrary threshold, i.e. the median. LZC of this binary sequence is then computed as outlined above. Here, this was computed for each channel and averaged to give a single value of $C^{TS}$ for each subject.

An advantage of microstate LZC over time series LZC is that microstate LZC accounts for multivariate patterns in the data. Therefore we also compare against Omega complexity, which is a widely used multivariate measure of EEG complexity [29,32–36] based on information theory, and uses information contained in the covariance matrix to estimate the complexity of the path of the EEG through $N$ dimensional space, where $N$ is the number of channels. Given EEG with a covariance matrix which has eigenvalues $\lambda_i$ (where $i = 1,...,N$ and $N$ is the number of EEG electrodes), Omega complexity ($\Omega$) is given by

$$\Omega = \exp\left(-\sum_{i=1}^{N} \frac{\lambda_i}{\Lambda} \cdot \log\left(\frac{\lambda_i}{\Lambda}\right)\right),$$

where, $\Lambda = \sum_{i=1}^{N} \lambda_i$. Omega complexity is close to zero if all channels follow a highly similar time course, whilst it is close to one if the channels follow very different time courses.

In this data, both classical measures demonstrated significant reductions in AD (Ω, $P$=.0191; $C^{TS}$, $P$=.0127; Figure 3), however the effect size of these differences were notably lower than for $C$, suggesting they are less sensitive to alterations to EEG complexity in AD. No significant correlation existed between $C$ and $C^{TS}$ ($r$ = 0.1363, $P$=.7222), or $C$ and Ω ($r$ = 0.3002, $P$=.0806). Omega complexity predominantly captures spatial complexity, being founded in covariance of time series, whilst time series LZC predominantly captures temporal complexity and is a univariate measure. The larger effect size of microstate LZC suggests that additional information can be gained by capturing both spatial and temporal patterns.

### 4. Use of eyes-open data

A key methodological factor that may affect results and therefore must be addressed is the use of eyes-open EEG, since the majority of past EEG microstate studies have been performed on eyes-closed resting state data.

A detailed study of differences in EEG microstates between eyes-open and eyes-closed conditions was performed by Seitzman et al. [37]. While the eyes-open data typically had less variance explained than the eyes-closed state, the curves of variance explained vs number of classes had almost identical forms suggesting that optimization criteria such as Krzanowski-Lai [18] (used here to choose the optimum number of microstates), which typically search for the 'elbow' in such a curve, would identify optima at the same point in both states. Furthermore, Seitzman et al. found that, given four classes, the eyes-open classes largely matched those of the four classical eyes-closed classes. Therefore, the results of Seitzman et al. support our results that in eyes open data, four microstate classes are optimum, and the topographies of these classes should largely match those reported in eyes closed data.

The microstates presented here are shorter than those typically seen in the literature, which are usually more than 80 ms [16]. von Wegner et al. [38] found that microstate duration is closely related to frequency of neuronal oscillations in the EEG, and Khanna et al. [20] predicted that factors such as eyes-open data and aging would result in reduced microstate durations due to different dominant frequencies. Comparisons of eyes-open and eyes-closed microstates further supports this, demonstrating reduced microstate durations in the eyes-open data [37,39]. Therefore, it is possible that an identical study in eyes-closed data may result in different durations to this study.

### 5. Supplementary Figures

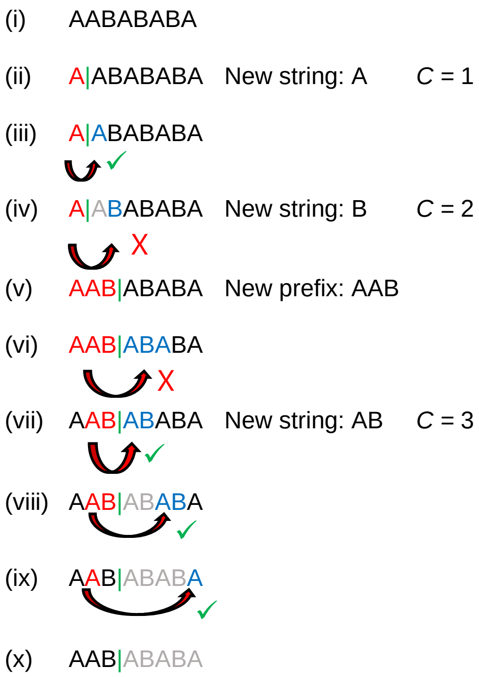


Figure S1: **Calculation of LZC from a string**. (i) We aim to calculate the LZC from a string *S* consisting of A's and B's. The *k*'th entry of *S* is denoted *S*[*k*]. (ii) We begin with a prefix of length one, demonstrated by the vertical green line separating the prefix from the rest of the string. The aim is to describe *S* in terms of a small number of substrings of the prefix (pointers). LZC (*C*) is the number of distinct substrings in the data. The initial prefix (A) is a new substring, so we initialize *C* = 1. (iii) Working from left to right, the pointer matches *S*[2] (both are the letter A), so complexity does not increase. (iv) However, *S*[3] is the letter B and hence does not match the pointer. Therefore a new substring has been identified (B), and *C* increases to 2. (v) Since the current prefix is not sufficient to describe the whole data, we then move it right to the point at which there is no match, making a new prefix AAB. (vi) Pointer AAB does not match *S*[4,5,6], so we next explore substrings of the prefix as the pointer. (vii) Pointer AB matches with *S*[4,5], so a new string is defined and complexity increases (*C* = 3). (viii) Furthermore the pointer AB also matches with *S*[6,7]. (ix) The pointer A matches with *S*[8]. (x) *S* is fully described by these 3 substrings, meaning the LZC of *S* is 3.

### 6. Supplementary Tables

| 1. SWE HOA | SWE AD | SWE MCIs | SWE MCIc | RSM HOA | RSM AD |
|---|---|---|---|---|---|
| SWE HOA | 0.1695 (0.3339) | 0.1781 (0.3339) | 0.6238 (0.6743) | 0.0338 (0.1015) | 0.1720 (0.3339) |
| SWE AD | | 0.9364 (0.9364) | 0.2346 (0.3911) | 0.0155 (0.1015) | 0.0233 (0.1015) |
| SWE MCIs | | | 0.2909 (0.4364) | 0.0338 (0.1015) | 0.0194 (0.1015) |
| SWE MCIc | | | | 0.4909 (0.6136) | 0.6294 (0.6743) |
| RSM HOA | | | | | 0.4191 (0.5714) |
| RSM AD | | | | | |

Table S1: ***p*-values for pairwise comparisons of the six cohorts for age.** Uncorrected *p*-values are reported, with false discovery rate corrected *p*-values in brackets accounting for multiple hypotheses.

| | SWE HOA | SWE AD | SWE MCIs | SWE MCIc |
|---|---|---|---|---|
| SWE HOA | | $1 \times 10^{-7}(7 \times 10^{-7})$ | $3 \times 10^{-4}(0.001)$ | 0.0138 (0.0207) |
| SWE AD | | | 0.0032(0.0065) | 0.0253 (0.0304) |
| SWE MCIs | | | | 0.7939 (0.7939) |
| SWE MCIc | | | | |

Table S2: **Uncorrected *p*-values for pairwise comparisons of the six cohorts for MMSE.** Uncorrected *p*-values are reported, with false discovery rate corrected *p*-values in brackets accounting for multiple hypotheses.

| Source | Sum Sq. | DF | Mean Sq. | F | P |
|---|---|---|---|---|---|
| Class | 237.2 | 3 | 79.1 | 0.67 | 0.5674 |
| Group | 1162.7 | 1 | 1162.7 | 9.95 | 0.0019 |
| Class*Group | 430.7 | 3 | 143.6 | 1.23 | 0.3007 |
| Error | 21029 | 180 | 116.8 | | |
| Total | 22809 | 187 | | | |

Table S3: Two-way ANOVA table for mean duration of microstates. The class term refers to the four microstate classes, whilst the group term refers to clinical diagnosis, i.e. AD vs HOA.

| Source | Sum Sq. | DF | Mean Sq. | F | P |
| --- | --- | --- | --- | --- | --- |
| Class | 0.0256 | 2 | 0.0128 | 1.72 | 0.1803 |
| Class*Group | 0.0040 | 2 | 0.0020 | 0.27 | 0.7657 |
| Error | 1.3478 | 182 | 0.0074 | | |
| Total | 1.3791 | 187 | | | |

Table S4: Two-way ANOVA table for coverage of microstate classes. The class term refers to the four microstate classes, whilst the group term refers to clinical diagnosis, i.e. AD vs HOA.

| Source | Sum Sq. | DF | Mean Sq. | F | P |
| --- | --- | --- | --- | --- | --- |
| Edge | 0.0037 | 6 | 6.1e-4 | 0.5891 | 0.7391 |
| Edge*Group | 0.0067 | 6 | 0.0011 | 1.0725 | 0.3777 |
| Error | 0.5688 | 546 | 0.0010 | | |
| Total | 0.5835 | 563 | | | |

Table S5: Two-way ANOVA table for Markovian switching between microstate classes. The edge term refers to the edges of the Markovian transition matrix, whilst the group term refers to clinical diagnosis, i.e. AD vs HOA.
